# The neocortical progenitor specification program is established through combined modulation of SHH and FGF signaling

**DOI:** 10.1101/841932

**Authors:** Odessa R. Yabut, Hui-Xuan Ng, Keejung Yoon, Jessica C. Arela, Thomas Ngo, Samuel J. Pleasure

## Abstract

Neuronal progenitors in the developing forebrain undergo dynamic competence states to ensure timely generation of specific excitatory and inhibitory neuronal subtypes from distinct neurogenic niches of the dorsal and ventral forebrain, respectively. Here we show evidence of progenitor plasticity when Sonic hedgehog (SHH) signaling is left unmodulated in the embryonic neocortex of the dorsal forebrain. At early stages of corticogenesis, loss of Suppressor of Fused (Sufu), a potent inhibitor of SHH signaling, in neocortical progenitors, altered their transcriptomic landscape. Ectopic activation of SHH signaling occurred, via degradation of Gli3R, resulting in significant upregulation of Fibroblast Growth Factor 15 (FGF15) gene expression. Consequently, activation of FGF signaling, and its downstream effector the MAPK signaling, facilitated expression of genes characteristic of ventral forebrain progenitors. Our studies identify the importance of modulating extrinsic niche signals such as SHH and FGF15 to maintain the competency and specification program of neocortical progenitors throughout corticogenesis.

**SIGNIFICANCE STATEMENT:** Low levels of FGF15 control progenitor proliferation and differentiation during neocortical development but little is known on how FGF15 expression is maintained. Our studies identified SHH signaling as a critical activator of FGF15 expression during corticogenesis. We found that Sufu, via Gli3R, ensured low levels of FGF15 was expressed to prevent abnormal specification of neocortical progenitors. These studies advance our knowledge on the molecular mechanisms guiding the generation of specific neocortical neuronal lineages, their implications in neurodevelopmental diseases, and may guide future studies on how progenitor cells may be utilized for brain repair.

## INTRODUCTION

The adult mammalian neocortex is composed of an intricate network of diverse excitatory and inhibitory neurons derived from distinct progenitor domains of the embryonic forebrain. Excitatory neurons originate from the ventricular and subventricular zones (VZ/SVZ) of the embryonic neocortex, while inhibitory neurons (interneurons) originate from the ganglionic eminences (GE). During corticogenesis, radial glial (RG) progenitors populating the VZ/SVZ sequentially generate deep-layer excitatory neurons, followed by upper-layer excitatory neurons via intermediate progenitor (IPC) or outer radial glial cells (Beattie and Hippenmeyer, 2017). This process must be tightly regulated since an imbalance between excitatory and inhibitory activity underlie a number of neurological and neuropsychiatric disorders (Sohal and Rubenstein, 2019).

A combination of intrinsic and extrinsic cues guides and maintains the specification program of neocortical progenitors throughout corticogenesis to generate neuronal diversity. But how these cues are integrated in neocortical progenitors to produce distinct neuronal subtypes in a temporal manner is still largely unclear. Our previous work identified fundamental mechanisms at early stages of corticogenesis ensuring proper specification of neocortical progenitors into distinct excitatory neuronal lineages, through modulation of Sonic hedgehog (SHH) signalling pathway (Yabut et al., 2015). SHH signalling is triggered upon binding of SHH ligands to the transmembrane receptor Patched 1 (Ptch1), which relieves its inhibition of another transmembrane protein, Smoothened (Smo). Consequently, Smo initiates a cascade of intracellular events promoting the nuclear translocation of Gli, a family transcription factors, to activate SHH target gene expression. However, intracellular checkpoints are present to modulate SHH signalling. In the developing neocortex, Suppressor of Fused (Sufu), a potent inhibitor of SHH signalling, is highly expressed in neocortical progenitors to modulate SHH signals to ensure the production of molecularly distinct upper and deep layer excitatory neurons (Yabut et al., 2015). Sufu exerted this effect by ensuring the stable formation of Gli transcription factors, the downstream effectors of SHH signalling. Specifically, loss of Sufu resulted in the degradation of the repressor form of Gli3 (Gli3R), the predominant Gli protein in the developing neocortex (Fotaki et al., 2006; Palma and Ruiz i Altaba, 2004; Wang et al., 2011; Wilson et al., 2012) leading to the production of misspecified neocortical progenitors by mid-corticogenesis. However, little is known on the identity of downstream molecular targets of SHH signalling or Gli3 in neocortical progenitors, and how deregulation of these targets due to uncontrolled SHH signalling might affect neocortical progenitor fates.

Here we show that endogenous levels of SHH, in the absence of Sufu, can sufficiently increase SHH signalling activity in neocortical progenitors resulting in drastic changes in the transcriptomic landscape of the VZ/SVZ at early stages of corticogenesis. In accordance to our previous findings, ventral forebrain progenitor gene transcripts are already ectopically expressed in neocortical progenitors in the embryonic (E) 12.5 neocortex mice lacking Sufu. Additionally, we find that activation of Fibroblast Growth Factor (FGF) signalling, via the upregulated gene expression of FGF15, leads to the misspecification of progenitors, particularly affecting the production of IPCs. These novel findings reveal how uncontrolled SHH signalling and its downstream gene targets can re-define progenitor competency in the embryonic neocortex. Further, this underscores the importance of intrinsic cellular responses, via modulatory proteins such as Sufu, to temporally restrain extrinsic niche signals that can influence progenitor identity and fate.

## RESULTS

### Specification defects are evident in discrete regions of the neocortex of E12.5 embryonic mice lacking Sufu

The role of SHH signaling in neocortical neuron specification is critical prior to E13.5, a timepoint at which superficial projection neurons are just beginning to differentiate. Analysis of mice in which Sufu is conditionally deleted at E10.5 in neocortical progenitors using the Emx1-Cre driver (*Emx1-cre/+;Sufu-fl/fl* or Sufu-cKO), revealed that modulating SHH signaling is critical to properly specify distinct superficial and deep layer projection neurons, after dorsoventral patterning of the forebrain (Yabut et al., 2015). While specification defects were clear at E14.5 in Sufu-cKO cortex, any molecular changes prior to this timepoint were not deeply examined in the previous study. Since changes in Gli2 and Gli3R levels were apparent at E12.5, we postulated that critical molecular alterations must have occurred at this timepoint. We therefore initiated our studies by careful examination of Pax6 expression, which is highly expressed in neocortical RG progenitors (Ypsilanti and Rubenstein, 2016). As expected, we found that Pax6 exclusively expressed in dorsal forebrain regions of the E12.5 control and Sufu-cKO brains, and not in the ganglionic eminence (GE) (**Figure 1A**). However, Pax6 expression was noticeably intermittent in anterior regions of the E12.5 Sufu-cKO neocortex (arrowheads in boxed regions, **Figure 1A**). Areas lacking Pax6 exhibited a columnar distribution hinting at anomalous RG clones (**Figure 1Ab**). Analysis of corresponding regions showed that the E14.5 Sufu-cKO neocortex similarly displayed columnar distribution of Pax6+ and Pax6-regions in anterior regions (**arrowheads, Figure 1B**), but this distribution was not prevalent in posterior regions (**Figure 1B**). These defects were not present at E10.5, in which the distribution of Pax6+ cells were largely indistinguishable between controls and Sufu-cKO embryos (**Figure 1-1**). Therefore, despite having properly formed dorsal forebrain domains, a subpopulation of neocortical RG progenitors displayed aberrant behavior in the E12.5 Sufu-cKO neocortex.

**Figure 1.**
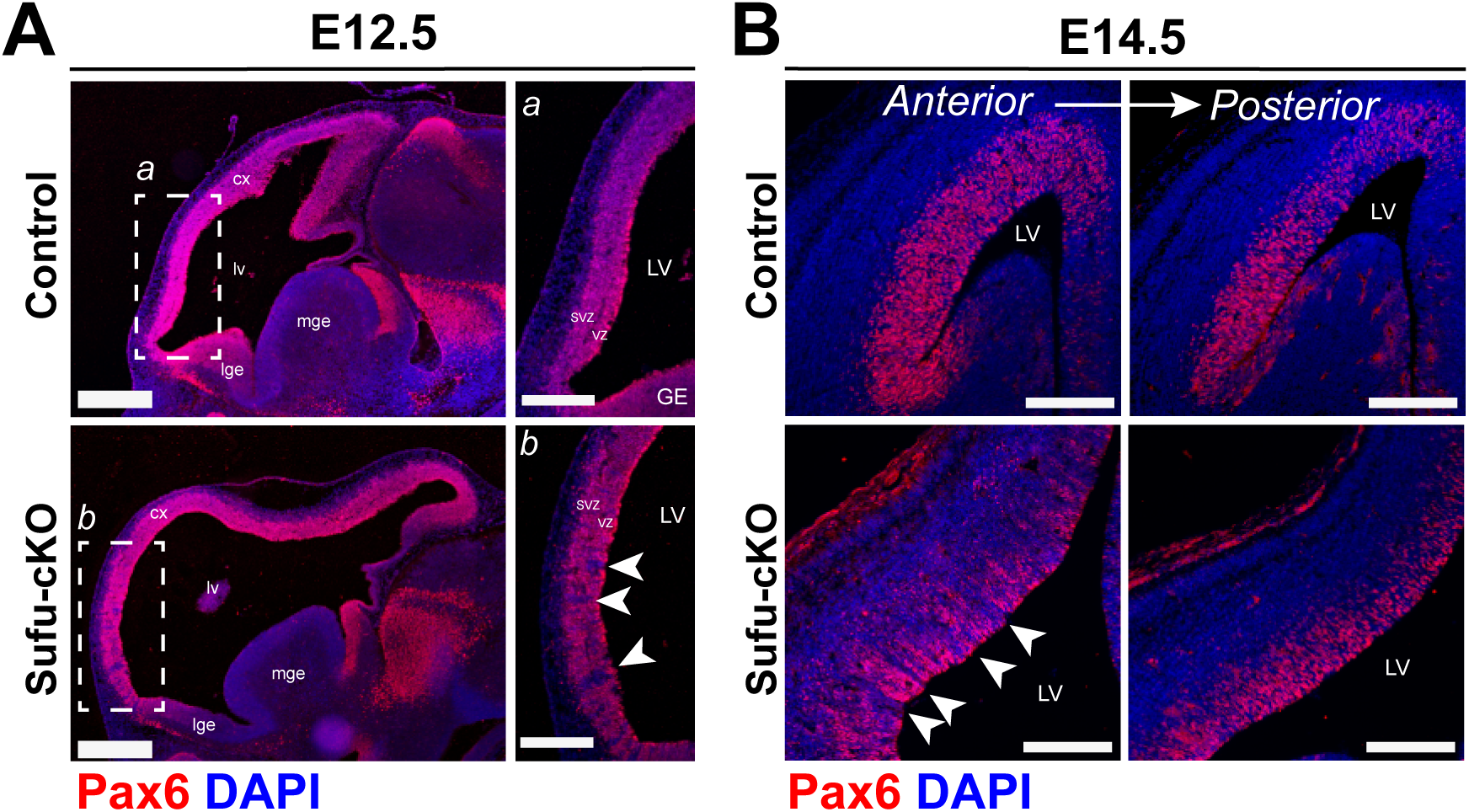
Neocortical progenitor defects are evident in discrete regions of the neocortex of the E12.5 mice lacking Sufu. **(A)** Immunofluorescence staining using dorsal forebrain progenitor marker, Pax6, and DAPI counterstain, show high Pax6 expression in the dorsal forebrain (cx) compared to the lateral (LGE) or medial (MGE) ganglionic eminence in both the E12.5 control and Sufu-cKO embryonic forebrains. Higher magnification of boxed regions in A and B show low or absent Pax6 expression in specific areas of the rostral neocortex of Sufu-cKO forebrains (arrows) but not in controls. Sections are counterstained with DAPI. Scale bars = 500 μm or 250 μm (boxed region). **(B)** Pax6 immunofluorescence staining show areas lacking Pax6 expression in the anterior neocortex of the E14.5 Sufu-cKO mice (arrows) but not in posterior regions or in controls. Scale bar = 200 μm.

### Upregulated expression of SHH signaling targets in Sufu mutant neocortical progenitors

To better understand the molecular changes in neocortical progenitors of the E12.5 Sufu-cKO neocortex, we isolated total RNA from dissected control and mutant dorsal forebrain for transcriptome profiling by RNA-Seq (**Figure 2A**). We confirmed that SHH signaling gene targets such as *Gli1*, *Patched 1* and *2* (*Ptch1 and Ptch2*), and the *Hedgehog-Interacting Protein* (*Hhip*) are specifically upregulated in the E12.5 Sufu-cKO neocortex compared to controls (**Figure 2B**). We validated these observations by *in situ* hybridization using probes for *Ptch1*, which was ectopically expressed throughout the neocortical expanse (**Figure 2E-2F**) in contrast to controls (**Figure 2C-2D**). Levels of *Ptch1* expression were confined within the VZ/SVZ across the cortical and hippocampal primordia (**Figure 2G-2H**) and were particularly high in rostral neocortical regions. Interestingly, expression of *Ptch1* also followed a visible columnar pattern (**Figure 2G-2H, arrows**) along the anterior neocortex of the E12.5 Sufu-cKO mice. These findings indicated deregulation of SHH signaling in discrete neocortical progenitor subpopulations, and not differentiated neurons, in the E12.5 neocortex of Sufu-cKO mice.

**Figure 2.**
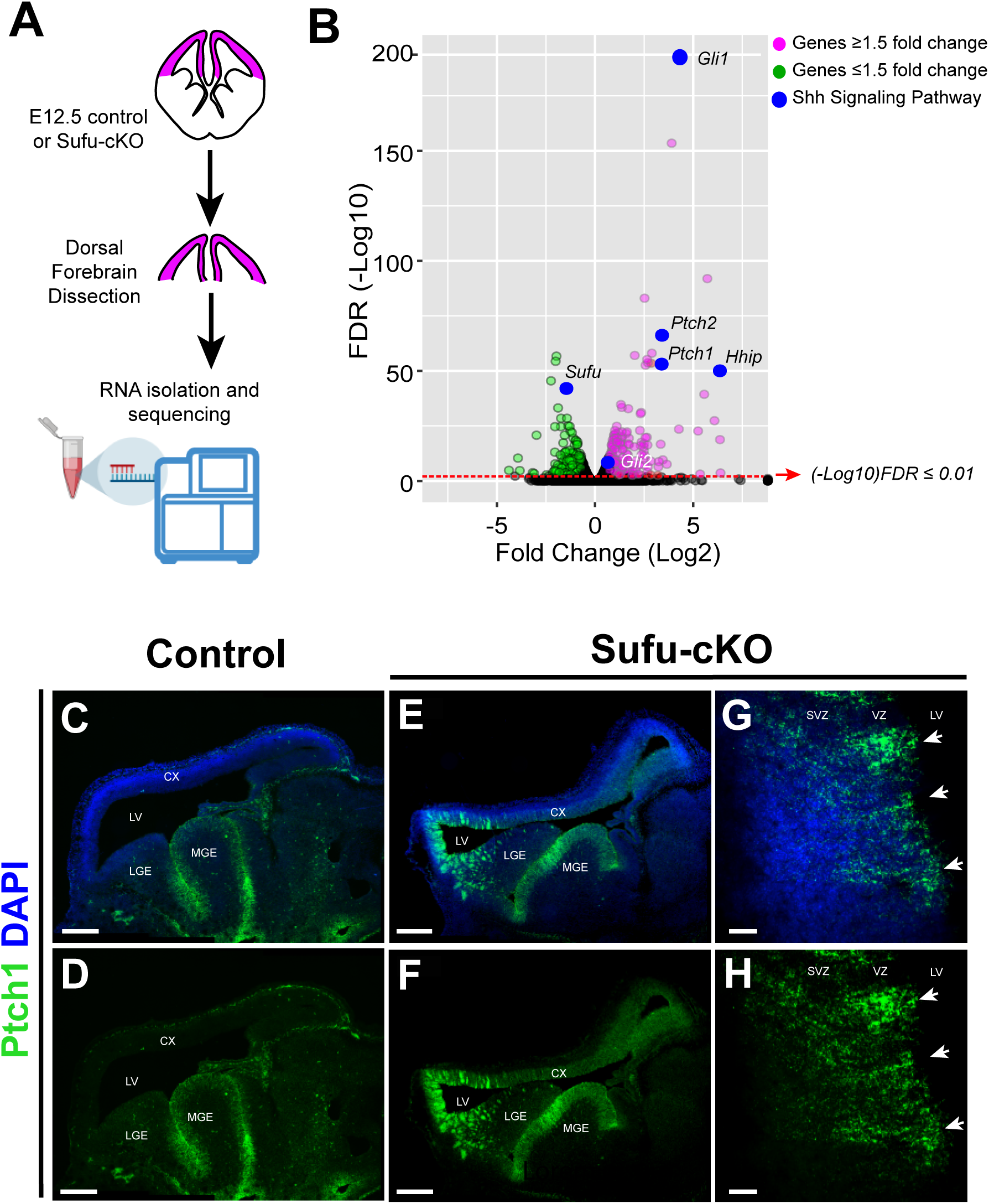
Upregulated expression of SHH signaling gene targets in neocortical progenitors of the E12.5 Sufu-cKO dorsal forebrain. **(A)** Schematic showing dorsal forebrain areas (pink) dissected from control and mutant E12.5 mice for RNA-Seq analysis. **(B)** Volcano plot of RNA-Seq data set highlighting differentially expressed genes with adjusted *p* value < 0.01 (FDR (-Log10)) and Fold Change (Log2) ≥ 1.5 (red circles) or Fold Change (Log2) ≤ 1.5 (green circles), and genes in the SHH signaling pathway (blue circles), between the E12.5 dorsal forebrain of controls and Sufu-cKO E12.5 embryos. **(C-F)** RNAscope *in situ* hybridization on sagittal brain sections using probes for Patched 1 (Ptch1) validates upregulation of Ptch1 RNA expression in the E12.5 Sufu-cKO dorsal forebrain (E, F) whereas Ptch1 RNA expression is only detected in the MGE of controls (C, D). Sections in C and E are counterstained with DAPI. Scale bar = 500 μm. **(G, H)** Higher magnification of rostral neocortex of E12.5 Sufu-cKO dorsal forebrain showing Ptch1 RNA expression is preferentially higher along the ventricular zone (VZ) and subventricular zone (SVZ) where neocortical progenitors are localized. Ptch1 expression also appear in columns, radiating inward from the apical VZ (arrows). Sections in G is counterstained with DAPI. Scale bar = 25 μm.

### Altered Molecular Identity of Progenitors in the E12.5 Sufu-cKO neocortex

Since changes in SHH signaling activity in the neocortex are known to disrupt progenitor fate specification in late-stage corticogenesis (Komada et al., 2008a; Wang et al., 2016), we wondered if the ectopic activation of SHH signaling at E12.5 has initiated a cascade of disruptive differentiation events. Enrichment analyses found over-representation of genes with gene ontology (GO) terms associated with neural development, commitment, specification and differentiation (**Figure 3A**). Further examination of specific gene expression showed relatively mild changes in the expression of genes typical of dorsal forebrain progenitors (**Figure 3B and Table 1**). Indeed, similar to Pax6 expression in **Figure 1**, other markers for dorsal forebrain cells such as *Tbr2, Lhx2,* and *Nr2f1*, remained expressed, and may be even expressed at slightly higher levels in the mutant neocortex as observed with *Pax6, Tbr1*, *Nr2f1* or *Nr2f2* (**Figure 3B and Table 1**). These findings validated the efficiency of the dissection and confirmed that the dorsal identity of neocortical progenitor domains was established in the E12.5 Sufu-cKO neocortex.

**Figure 3.**
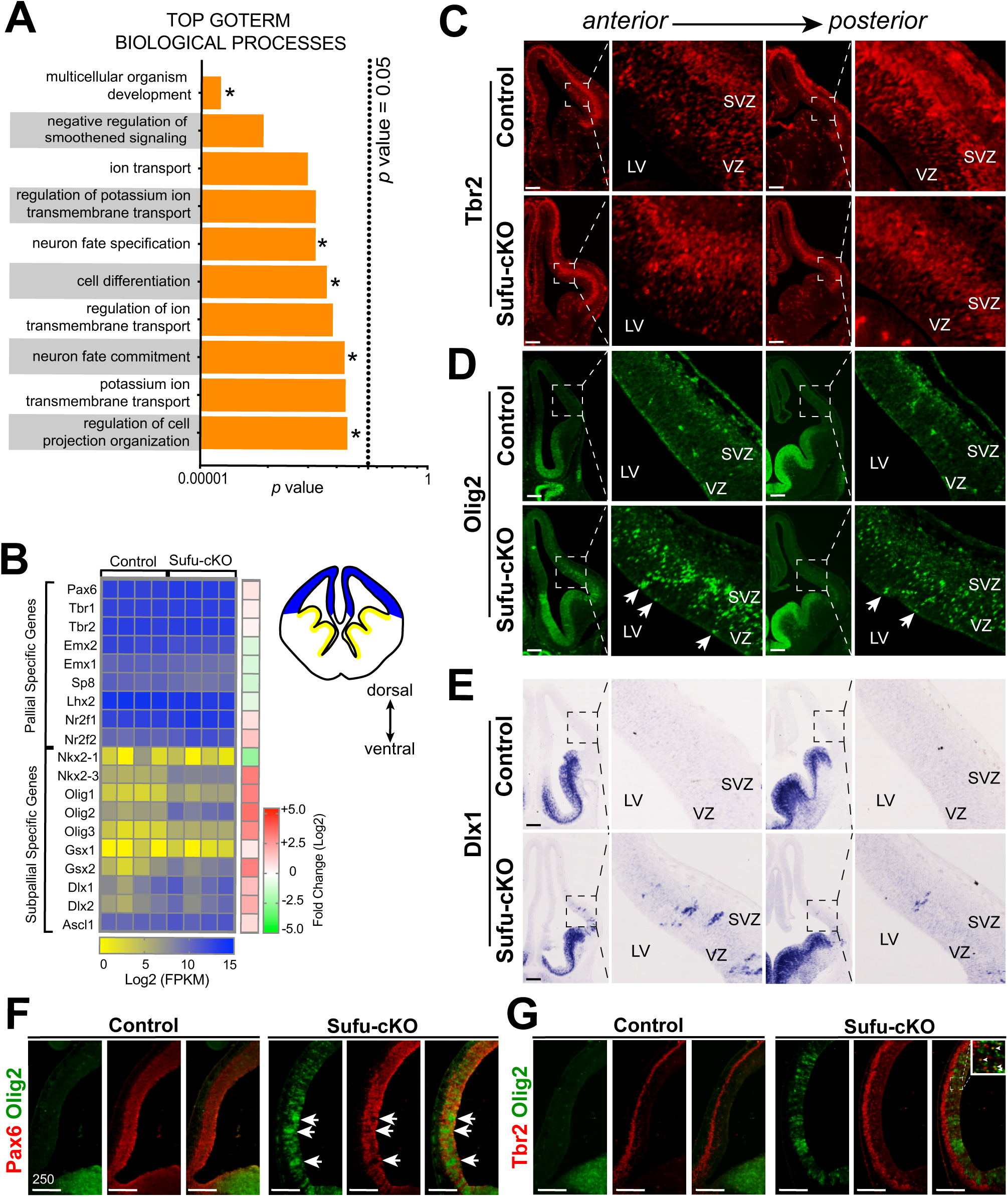
Increased expression of ventral progenitor markers in neocortical progenitors of E12.5 Sufu-cKO embryos. **(A)** Functional annotation of differentially expressed genes identified by RNA-Seq show top GOTERM Biological Processes (with adjusted *p* value < 0.05) involve development, specification, differentiation, and fate commitment (asterisks). There is also a notable enrichment in ion transmembrane transport GOTERMs reflecting disrupted electrophysiological properties due to abnormal differentiation of neurons or specific neuronal subtypes. **(B)** Heat map of select genes typically expressed by dorsal or ventral progenitors in individual control and Sufu-cKO mice (n=4 mice per genotype). RNA levels (Log2 FPKM scale) reflect mild differences in expression of dorsal progenitor genes (reflected by Fold Change scale), while dramatic differences in expression levels of ventral progenitor genes is observed between controls and Sufu-cKO dorsal forebrain. See also **Table 1**. **(C)** Immunofluorescence staining for Tbr2 in coronal sections of the E12.5 Sufu-cKO and control forebrain showed exclusive expression in the neocortex across the A-P axis. Scale bar = 100 μm. **(D-E)** Immunofluorescence staining for Olig2 (D) and in situ hybridization for Dlx1 on coronal sections of the E12.5 control and Sufu-cKO forebrain. These experiments validate the ectopic RNA expression of subpallial-specific genes across the A-P axis of the E12.5 Sufu-cKO neocortex whereas these genes were absent in controls. Olig2- and Dlx1-expressing cells largely localized in the VZ and SVZ. Some groups of cells expressing Olig2 and Dlx1 also appeared in columnar arrangement (arrows). Scale bar = 100 μm. **(H)** Double immunofluorescence staining on E12.5 sagittal sections with Pax6 and Olig2, a ventral forebrain progenitor marker, showed ectopic expression of Olig2 in areas where Pax6 is missing in the Sufu-cKO neocortex (arrows), whereas Olig2 was not expressed in this region in the control neocortex. Scale bar = 250 μm. **(I)** Double immunofluorescence staining with Tbr2 and Olig2 on sagittal sections of E12 Sufu-cKO and control littermates, showed that unlike Pax6+ cells, the distribution of Tbr2+ cells were not affected in the anterior regions where ectopic expression of Olig2 was present. However, Tbr2+ cells were found to co-express Olig2 in more dorsal regions of the E12.5 Sufu-cKO neocortex. Scale bar = 250 μm.

Nevertheless, RNA levels for several ventral progenitor genes dramatically increased in the E12.5 Sufu-cKO neocortex compared to controls (**Figure 3B**). We found a specific increase of subpallial-specific genes in the neocortex (**Figure 3B and Table 1**). Moreover, while we previously did not observe a significant increase in Ascl1 protein expression in the E12.5 neocortex (Yabut et al., 2015), here we found significantly higher levels of Ascl1 transcript, despite not detecting Ascl1 protein (**Extended Data Figure 3-1A**). Additionally, significant upregulation of genes normally expressed in the GE such as *Gsx2,* and *Dlx1/2* (Petryniak et al., 2007) were also ectopically expressed in the neocortex of E12.5 Sufu-cKO mice (**Figure 3B**).

We subsequently conducted immunostaining or *in situ* hybridization experiments to validate the expression of subpallial-specific markers. In agreement with the transcript increase quantified by RNA-Seq, visibly higher levels of Tbr2+, NR2F1+, and Lhx2+ cells were observed across the anterior to posterior (AP) axis of the Sufu-cKO neocortex compared to controls, (**Figure 3B-3C**, **Extended Data Figure 3-1B** to **3-1C**). Similarly, ectopic expression of subpallial-specific genes Olig2, Dlx1, and Gsx1 were detected in the E12.5 Sufu-cKO neocortex (**Figure 3D-3E, Extended Data Figure 3-1D**). Expression of these genes were detected in the SVZ and VZ regions of the E12.5 Sufu-cKO neocortex and exhibited a columnar pattern, whereas these genes were absent in controls. Additionally, neocortical progenitors in these regions were improperly specified since we detected ectopic expression of the ventral forebrain progenitor marker, Olig2, in areas where Pax6 was absent in the E12.5 Sufu-cKO neocortex (**arrows in Figure 3F**), whereas Olig2 was completely absent in the neocortex of control mice. This expression pattern persisted in the anterior regions of the E14.5 Sufu-cKO neocortex but not in posterior regions (**Extended Data Figure 3-2A**). However, we did not see similarly extensive disruptions in Tbr2 expression in the E12.5 Sufu-cKO neocortex, even in areas where Olig2+ cells were highly enriched (**Figure 3C and 3G**). Nevertheless, while the majority of Tbr2+ cells in the E12.5 Sufu-cKO SVZ did not co-express Olig2, a few cells within the VZ co-expressed Olig2 and Tbr2 (arrowheads **boxed inset, Figure 3G**). Further, by E14.5, Tbr2+ cells, similar to Pax6+ cells, became intermittent in the anterior neocortex of Sufu-cKO mice and were populated by Ascl1+ cells (**Extended Data Figure 3-2B, 3-2C**). Ectopic expression of Olig2 was not prevalent in the E11.5 Sufu-cKO neocortex although we noted irregularities in Olig2 expression near the pallial-subpallial boundary (**Extended Data Figure 3-1E**), indicating that a subset of aberrant progenitors may be present at this stage. Altogether, these findings establish that activation of SHH signaling in early stages of corticogenesis did not disrupt the regionalization of dorsoventral axis but has begun to destabilize the specification program of neocortical RG progenitors to disrupt the specification of Tbr2+ IP cells.

### Ectopic activation of SHH signaling upregulates FGF15 expression

To determine how these genetic changes mediated specification defects in the E12.5 Sufu-cKO neocortex, we further analyzed the overall nature of differentially expressed genes from our RNA-Seq data (Table 2). Functional analysis of the transcriptome using DAVID (https://david.ncifcrf.gov/) showed enrichment of genes encoding proteins with roles in cell-cell communications such as membrane-bound or extracellular matrix proteins in the E12.5 Sufu-cKO transcriptome (**Figure 4A**). Thus, the molecular make-up of the VZ/SVZ progenitor niche has been significantly altered in response to the ectopic activation of SHH signaling in the E12.5 Sufu-cKO neocortex. Among these is the gene encoding the secreted ligand, *Fibroblast Growth Factor 15* (*Fgf15*) (**Figure 4B**). *Fgf15* was a previously reported SHH signaling gene target in the developing cerebellum affecting neuronal precursor behavior (Gimeno and Martinez, 2007; Kim et al., 2018; Komada et al., 2008b). Similarly, we found that in the E12.5 Sufu-cKO neocortex, *Fgf15* dramatically increased (+5.72 Log2 Fold Change, p-value < 0.0001), likely as a consequence of Gli3R loss. *In situ* hybridization using *Fgf15* riboprobes confirmed these findings, with *Fgf15* ectopically expressed throughout the neocortical wall of the E12.5 and E14.5 Sufu-cKO mice while *Fgf15* expression was relatively low in controls (**Figure 4C-4D**). We also observed upregulation of *Fgf15* in embryos in which Smo was constitutively active in neocortical progenitors (*Emx1-Cre;SmoM2* or SmoM2-cA) (Long et al., 2001), confirming the role of activated SHH signaling in inducing *Fgf15* gene expression in the embryonic neocortex (**Extended Data 4-1**). Importantly, *Fgf15* expression in the E12.5 Sufu-cKO was detected along the VZ/SVZ, and particularly overlapped with *Ptch1*-expressing cells in the VZ (**Figure 4E**). These observations indicated that ectopic *Fgf15* expression was induced in RG progenitors along the VZ and persisted in intermediate progenitors as a consequence of loss of Sufu and deregulated SHH signaling.

**Figure 4.**
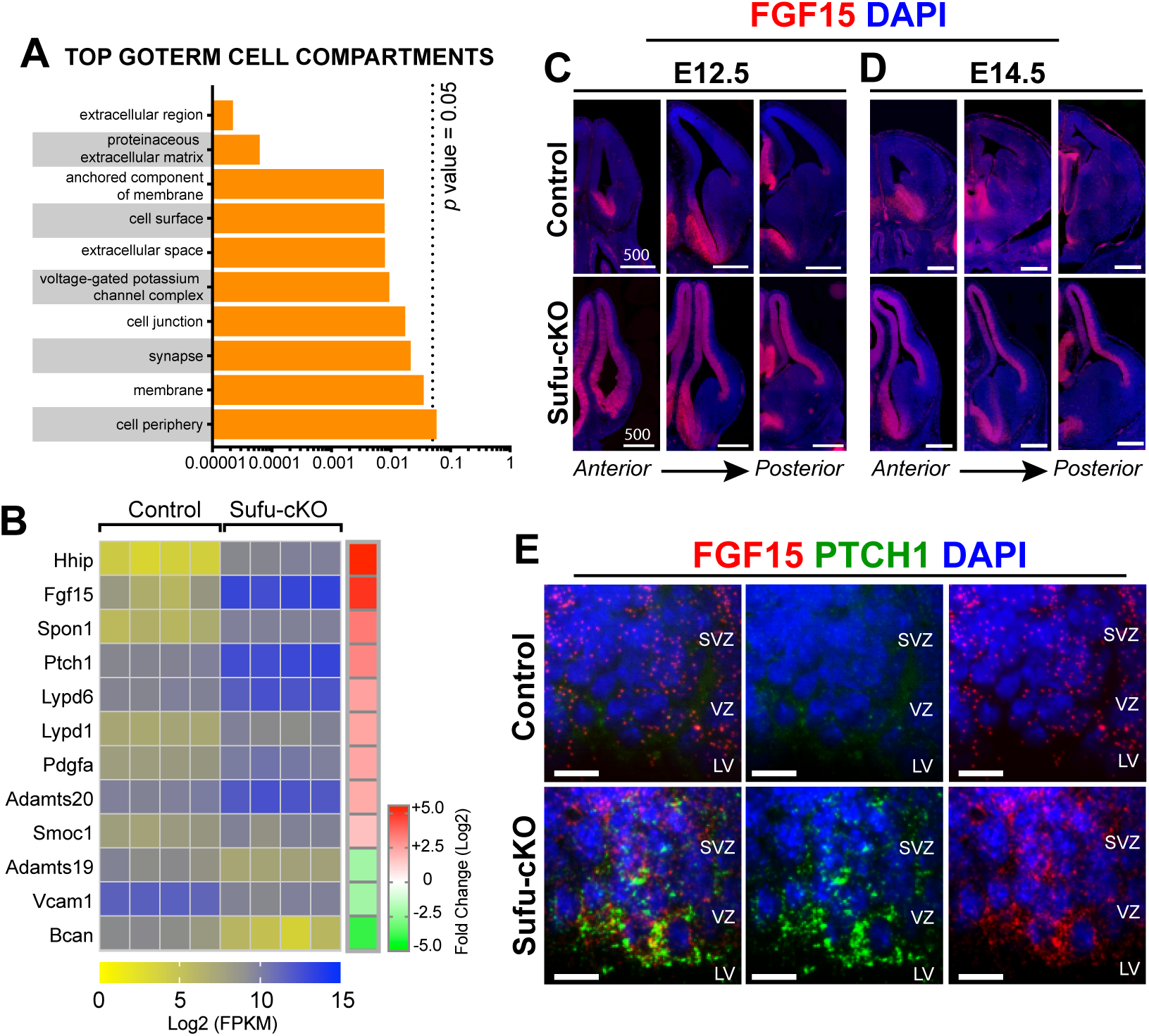
Ectopic activation of SHH signaling drives Fgf15 expression in neocortical progenitors of E12.5 Sufu-cKO embryos. **(A)** Functional annotation of differentially expressed genes identified by RNA-Seq showing the majority of genes encode proteins that localize to extracellular matrix or cell surface/membrane as the top GOTERMs Cell Compartments (with adjusted *p* value < 0.05). See also **Table 2**. **(B)** Heat map of top differentially expressed genes encoding extracellular matrix or cell membrane bound proteins between control and Sufu-cKO mice (n=4 mice per genotype). RNA levels (Log2 FPKM scale) show expression of *Fgf15* is significantly upregulated (reflected by Fold Change scale in the E12.5 Sufu-cKO dorsal forebrain. See also **Table 3**. **(C, D).** *In situ* hybridization for *Fgf15* in the E12.5 and E14.5 control and Sufu-cKO neocortex. High levels of *Fgf15* expression was detected throughout the E12.5 Sufu-cKO dorsal forebrain, and particularly enriched in the VZ/SVZ regions, whereas *Fgf15* expression was detected only in ventral forebrain regions in controls (C). Expression of *Fgf15* persisted in the E14.5 control and Sufu-cKO forebrains (D). This confirmed that activation of SHH signaling and loss of Sufu force Fgf15 expression in the embryonic neocortex. Scale bar = 500 μm. **(E)** Multiplex RNAscope ISH of Ptch1 and Fgf15 riboprobes on E12.5 brains did not detect Ptch1 expression, while low levels of Fgf15 expression was detected in the VZ and SVZ of the neocortex of controls. In the E12.5 Sufu-cKO neocortex, high levels of Ptch1 and Fgf15 colocalization was detected in the VZ and SVZ. Sections are counterstained with DAPI. Scale bar = 10 μm.

### Upregulated FGF15 expression correlates with ectopic activation of MAPK signaling in neocortical progenitor zones

FGF15 preferentially binds to its cognate receptor, FGF receptor 4 (FGFR4), to activate intracellular signaling cascades, particularly the Ras/mitogen activated protein kinase (MAPK) pathway (Guillemot and Zimmer, 2011). Indeed, in the neocortex of E12.5 Sufu-cKO mice, MAPK signaling pathway activity, as marked by phosphorylated-ERK1/2 (pERK1/2+) labeling, is visibly upregulated unlike controls (**Figure 5A**). We found that pERK1/2+ areas largely occupied the progenitor-rich VZ/SVZ neocortical regions whereas it was largely undetected in similar neocortical regions in controls. Quantification of pERK1/2+ regions in representative sections across dorsal forebrain (**Figure 5C**) showed a consistently larger areas with pERK1/2+ immunoreactivity in the E12.5 Sufu-cKO compared to controls (**Figure 5D**). This remained true at E14.5, where pERK1/2+-rich regions were detected further towards the dorsal regions of the Sufu-cKO neocortex (**Figure 5B**). Quantification of pERK1/2+ regions in the E14.5 neocortex confirmed these observations and showed a significant increase in Sufu-cKO mice (**Figure 5D**). At both E12.5 and E14.5 timepoints, cells labeled with pERK1/2+ clearly overlapped with FGF15-expressing VZ/SVZ areas in the Sufu-cKO neocortex (**Figure 4D**). Taken together, these observations indicated that loss of Sufu resulted in the overexpression of FGF15 in the neocortex, subsequently driving the ectopic activation of FGF signaling to activate intracellular MAPK signaling in neocortical progenitors.

**Figure 5.**
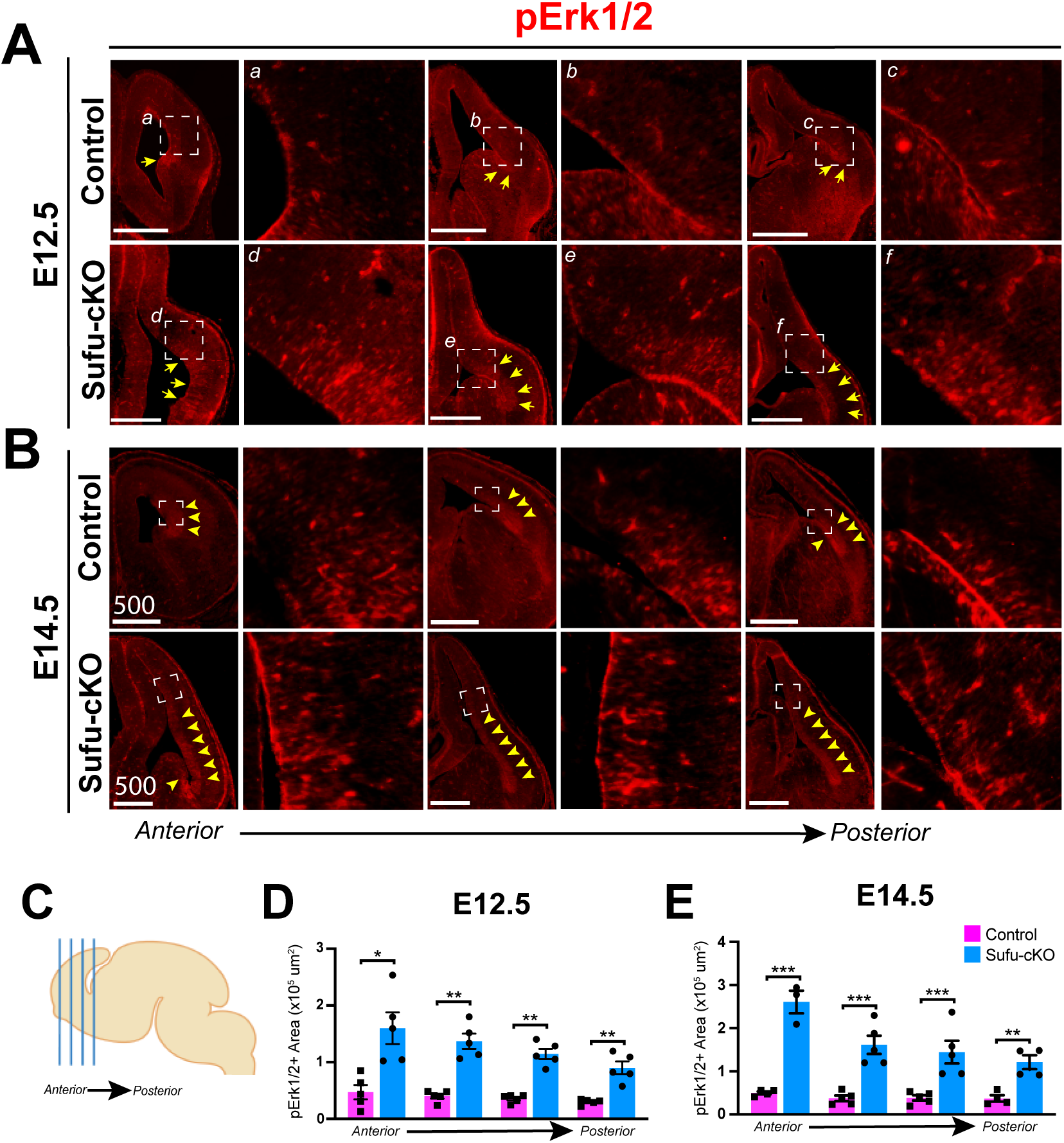
Upregulated FGF15 expression correlates with ectopic activation of MAPK signaling in neocortical progenitor zones. **(A, B)** Immunofluorescence staining against Phosphorylated Erk1/2 (pErk1/2) was conducted on E12.5 and E14.5 control and Sufu-cKO brains to detect for MAPK signaling activation. pErk1/2-expressing cells (pErk1/2+) were detected along the pallial-subpallial boundary (PSB) of the E12.5 control and Sufu-cKO forebrain (white arrows). However, pErk1/2+ cells were found from the PSB and the lateral cortex, particularly in the VZ/SVZ regions (yellow arrows, A). At E14.5, pErk1/2+ cells expanded dorsally within the VZ/SVZ regions in both control and Sufu-cKO neocortex (B). However, pErk1/2+ cells in the Sufu-cKO neocortex greatly expanded compared to controls (yellow arrows). Scale bars = 100 μm (A, B) and 200 μm. **(C-E)** Four representative sections across the A-P axis of the forebrain (C) were sampled to measure pERK1/2+ regions in the E12.5 and E14.5 control and Sufu-cKO mice. Bar graphs of quantification of pErk1/2+ regions in the neocortex of E12.5 (D) and E14.5 (E) control and Sufu-cKO mice (n=3-5 embryos per genotype) showing significant interaction between position and genotype (RM 2-way ANOVA, *p value* = 0.0365). Asterisks show significance between genotypes in Ph-Erk1/2+ regions in the Sufu-cKO neocortex at both E12.5 and E14.5, particularly in anterior regions (\**p* value ≤ 0.05; \*\**p* value ≤ 0.01, ***p value ≤0.001, RM, 2-way ANOVA with Holm-Sidak’s multiple comparisons test).

### FGF15 upregulation is required to elicit Shh signaling-mediated defects in the production and specification of intermediate progenitors

Reduction in intermediate progenitors (IP) is a consistent phenotype in the embryonic neocortex of mice with excessive levels of SHH signaling, including Sufu-cKO mice (Dave et al., 2011; Komada et al., 2008a; Yabut et al., 2015). We therefore investigated whether downregulation of IPs in the neocortex due to ectopic SHH signaling is directly mediated by FGF15 signaling. To test this, we cultured wildtype forebrain slices from the anterior regions of E12.5 control and Sufu-cKO embryos (**Figure 6A**). Forebrain organotypic cultures maintain the three-dimensional structure of the VZ/SVZ niche, allowing for careful examination of how precisely added compounds affect progenitor behavior over time. Forebrain slices cultured for 2 days *in vitro* (DIV) maintain their anatomical features with well-preserved dorsal and ventral domains. Neocortical IPCs typically expressing Tbr2 (Hevner, 2019) were exclusively observed in the dorsal forebrain whereas ventral forebrain progenitors typically expressing Olig2 (Miyoshi et al., 2007) (**Figure 6B**). Addition of various compounds altered IPC numbers in neocortical regions of forebrain slices (**Figure 6C and 6D**). SHH ligands significantly decreased the number of Tbr2+ cells in neocortical slices after 2 DIV when compared to mock-treated controls. Similarly, Tbr2+ IPCs were significantly reduced upon addition of FGF15 alone or with SHH. However, addition of cyclopamine, which inhibits SHH signaling by rendering Smo inactive, did not alter the number of Tbr2+ IPCs. Instead, addition of cyclopamine and FGF15 significantly reduced the number of IPCs after 2 DIV (**Figure 6C and 6D**). These findings indicated that blocking transmembrane proteins upstream of the SHH signaling pathway cannot sufficiently alter IPC production. Additionally, expression of downstream SHH gene targets, particularly FGF15, is required to exert changes in neocortical IPCs of the developing neocortex.

**Figure 6.**
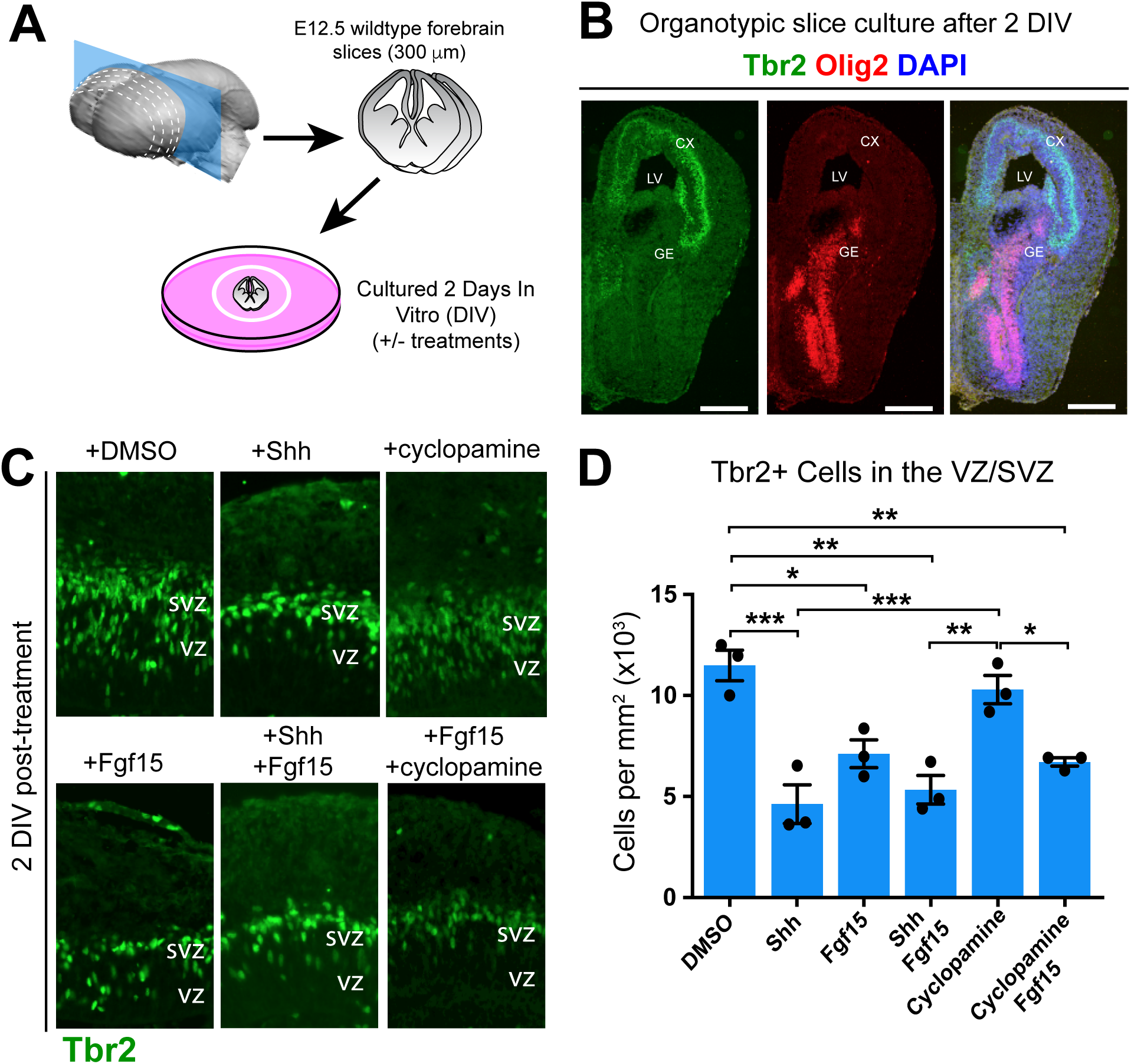
SHH signaling activate FGF15 signaling to inhibit production of neocortical intermediate progenitor cells. **(A)** Diagram of experimental design for organotypic forebrain slice cultures from wildtype E12.5 brains. **(B-D)** Immunofluorescence staining for dorsal (Tbr2, green) and ventral (Olig2, red) forebrain markers show exclusive localization of Tbr2 (B) and Olig2 (C)-expressing cells in dorsal and ventral forebrain regions, respectively. Merged images in (D) show no overlap in Tbr2 or Olig2 labeling. Scale bar = 500 μm **(E-K)** Immunofluorescence staining with Tbr2, an intermediate progenitor cell (IPC) marker of sectioned organotypic slice cultures fixed after 2 days in vitro (DIV). Slices treated with 200 ng/ml SHH (F) and 100 ng/ml FGF15 (H) show reduced numbers of Tbr2+ IPCs compared to slices treated with DMSO (E) or 5μM Cyclopamine (G). Combined FGF15 and SHH (I) or FGF15 and cyclopamine (J) also show reduced Tbr2+ IPCs. Quantification of Tbr2+ cells per unit area (K) confirm significant interaction between treatments (RM 2-way ANOVA, *p value* = 0.0001). Significant differences in Tbr2+ IPCs in SHH and FGF15-treated slice cultures (n = 3 experiments (2-3 slices each experiment) per treatment condition). Asterisks show significance between treatment conditions (\**p* value ≤ 0.05; \*\**p* value ≤ 0.01, ***p value ≤0.001, RM, one-way ANOVA with Holm-Sidak’s multiple comparisons test).

### High levels of FGF15 alter the specification program of neocortical progenitors

Ectopic SHH signaling in the developing neocortex ultimately results in the production of confused progenitors unable to maintain a specified neocortical neural fate (Yabut et al., 2015). The expansive ectopic activation of MAPK signaling, capable of altering neocortical progenitor fate (Wang et al., 2012), in the Sufu-cKO embryonic neocortex is a likely consequence of increasing levels of FGF15. We tested this by adding FGF15 in organotypic forebrain cultures and examined if this alone altered the fate of neocortical progenitors based on Olig2 expression. Indeed, we found that after 2 DIV, the decrease in Tbr2+ IPCs correlated with an obvious increase in Olig2+ cells in FGF15-treated slices compared to DMSO-treated controls (**Figure 7A-7D**). Further, low levels of Olig2 expression were detectable in the VZ region, where Olig2 is typically not expressed, and may co-express low levels of Tbr2, indicating that treatment of FGF15 alters the identity of RG progenitors transitioning into IPCs (**Figure 7G-7H, arrows**). Indeed, many Olig2+ cells in the SVZ also expressed Tbr2 in FGF15-treated slices compared to DMSO-treated controls (**Figure 7E, 7F**). Our quantification confirmed these observations, showing the approximately 4.5-fold increase in misspecified Tbr2+ cortical progenitors in FGF15-treated slices compared to DMSO-treated controls (**Figure 7I**). These findings indicate that excessive levels of FGF15 can sufficiently alter the identity of neocortical progenitors, leading to the failure to maintain a proper specification program in the developing neocortex.

**Figure 7.**
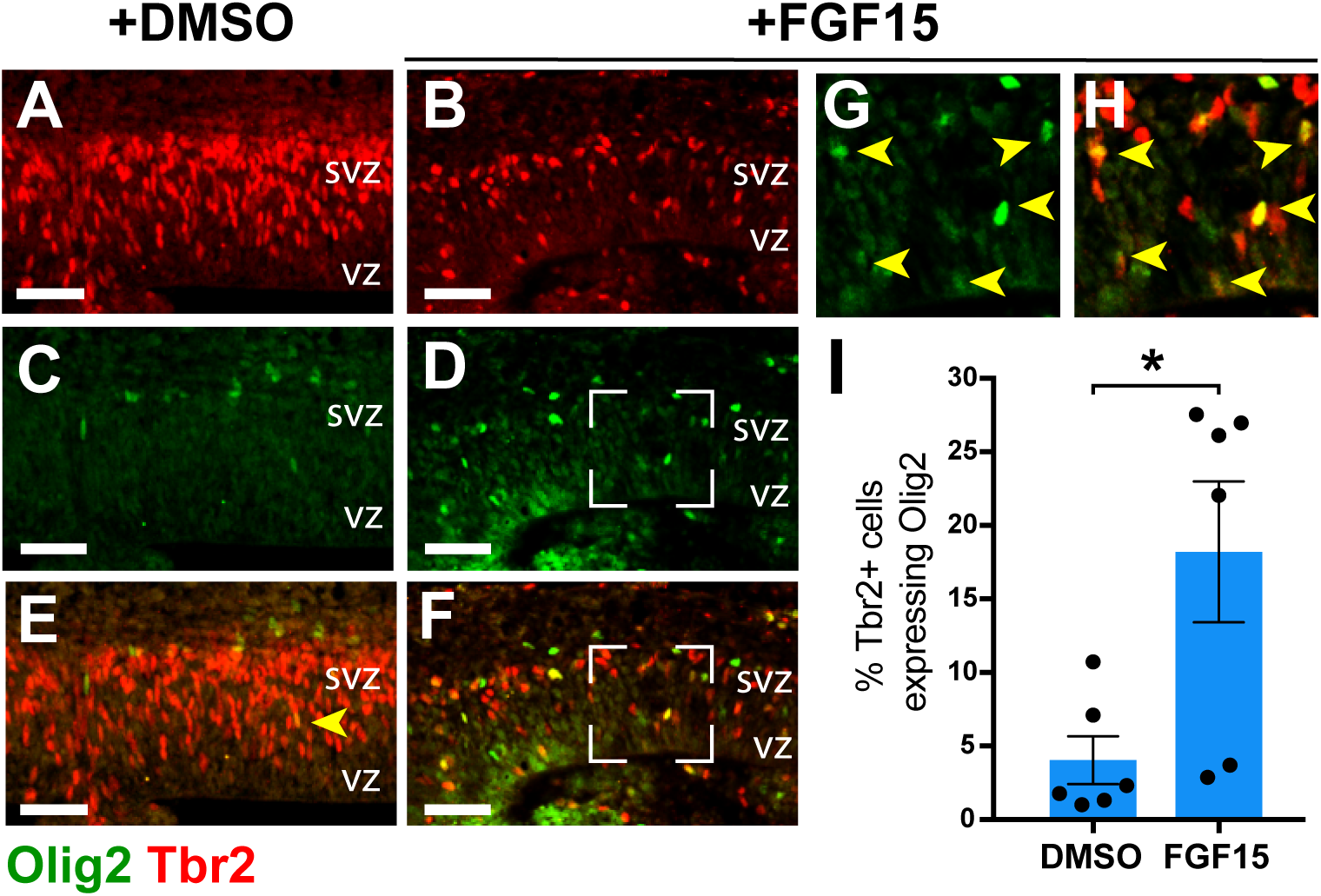
Increasing FGF15 levels alter the specification program of neocortical progenitors. (A-F) Double immunofluorescence staining with IPC marker Tbr2 (red), and the ventral progenitor marker Olig2 (green) on organotypic slice cultures fixed 2 DIV and post-treatment. DMSO-treated slices show an abundance of Tbr2+ IPCs in the SVZ (A) and some Olig2-expressing cells outside of the VZ/SVZ area (B). In contrast, Tbr2+ IPCs in FGF15-treated slices are fewer (B) and more Olig2+ cells are present in the VZ/SVZ (F). Although Tbr2+ cells expressing Olig2 were also sometimes observed in DMSO-treated slices (yellow arrow, C), the amount of double-labeled in FGF15-treated slices were visibly higher in the VZ/SVZ (yellow arrows, G and H). Scale bar = 50 μm (G) Graph represents the % of Tbr2+ cells colabeled with Olig2 in the VZ/SVZ of DMSO- and FGF15-treated slices (n = 3 experiments (2-3 slices each experiment) per treatment condition). The percentage of Tbr2+ co-expressing Olig2 is ∼4-fold in FGF15-treated over control DMSO-treated slices and is significantly higher (* = *p* value ≤ 0.05).

## DISCUSSION

Excitatory neurons in the mammalian neocortex are generated in a limited period at embryonic stages and mature into molecularly diverse subpopulations at postnatal stages. A strict specification program is maintained by neural progenitors to generate precise numbers and lineages, relying on spatially and temporally modulated molecular cues present in neurogenic niches of the embryonic forebrain. Our study identified SHH and FGF15 signaling as key pathways that must be tightly modulated to ensure successful differentiation of neocortical progenitors into distinct excitatory neuron lineages in the course of corticogenesis (**Figure 8**). These findings further underscore the importance of key modulatory factors at crucial timepoints in establishing and maintaining neocortical progenitor programs.

**Figure 8.**
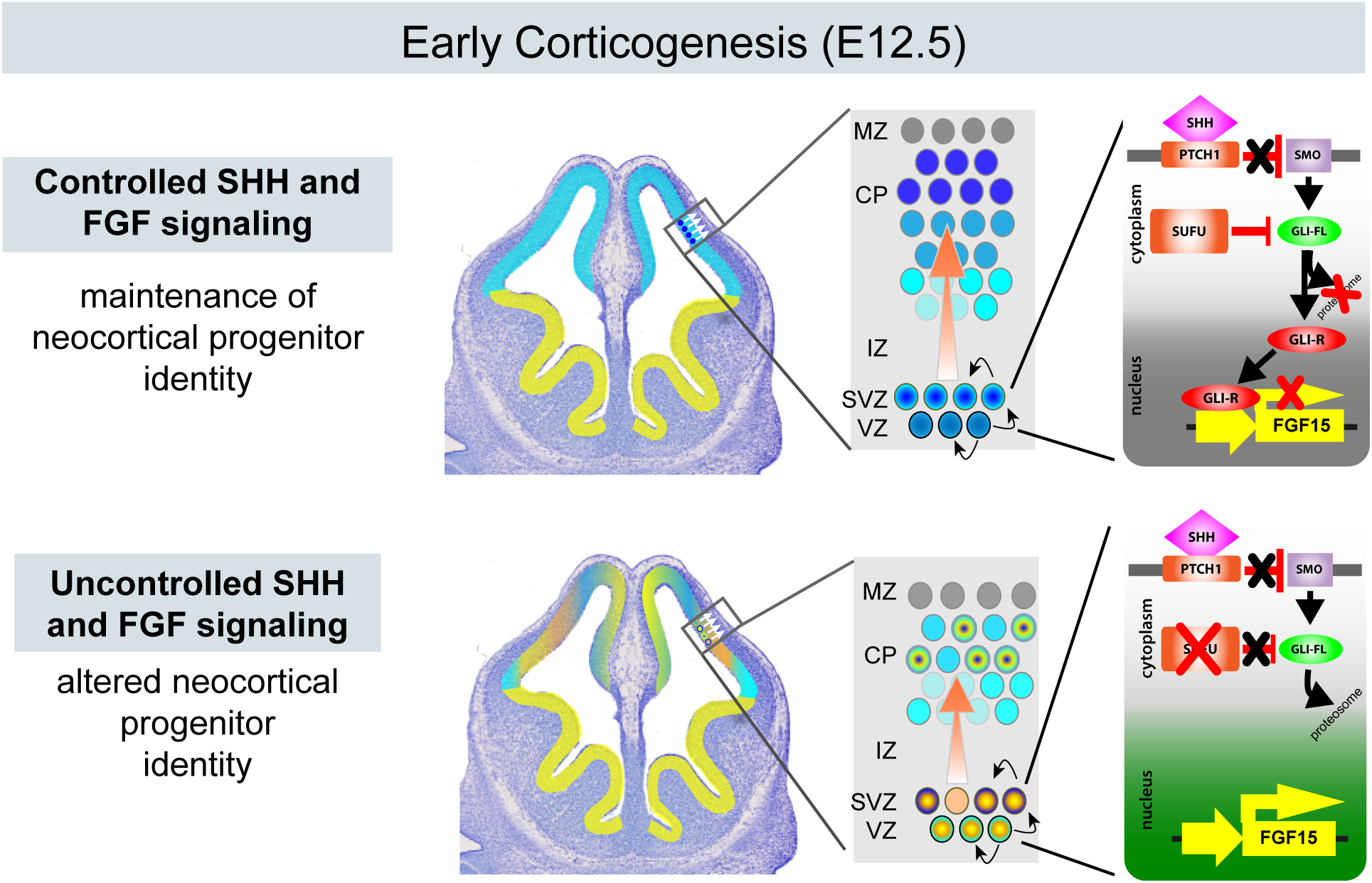
Schematic diagram of how modulation of SHH and FGF signaling affect neocortical progenitors at early stages of corticogenesis. Controlled SHH and FGF signaling, due to the repressive function of Gli3R, in the dorsal forebrain, maintains the neocortical identity and specification program of progenitors. Uncontrolled SHH and FGF signaling, achieved when Sufu expression is lost consequently leading to Gli3 degradation and the eventual upregulation of FGF15 expression, results in the inability of neocortical progenitors to maintain the dorsal forebrain identity and specification program.

### Neuronal lineage progression relies on the molecular makeup of the neurogenic niche at E12.5

Embryonic neocortical progenitors must follow strict lineage programs throughout corticogenesis to ensure the production of molecularly and functionally diverse excitatory neurons in the mature neocortex. Here, we determined that molecular events in early corticogenesis ensure proper lineage progression of neocortical progenitors. This process relies on tightly inhibiting Shh signaling activity. While pallial-specific progenitors, such as Pax6+ RGCs or Tbr2+ IPCs, are produced in the Sufu-cKO neocortex, these progenitors possessed underlying transcriptomic changes compromising their lineage progression. Many of these changes involved extracellular or plasma membrane bound proteins, which can substantially alter intercellular and extracellular interactions within the neurogenic niche. Further, we found that subpallial-specific gene transcripts were already ectopically expressed in the E12.5 Sufu-cKO. These indicate that while proteins encoded by these genes may not be detected in neocortical progenitors at E12.5, the stage has been set to alter their fates. Ultimately, these early alterations permanently deviate the lineage programs of neocortical progenitors, resulting in the production of misspecified excitatory neurons in the postnatal neocortex of Sufu-cKO mice.

### SHH signaling alters IPC production by preventing lineage progression of RGCs

IPC production is especially vulnerable to varying activity levels of SHH signaling. High SHH signaling activity inhibits IPC production at early stages of corticogenesis but promotes IPC production at later stages (Shikata et al., 2011; Wang et al., 2011, 2016; Yabut et al., 2016, 2015). While these studies established the mitogenic effect of Shh signaling to control neocortical progenitor proliferation, including IPCs, a growing number of studies are beginning to identify a role for SHH signaling in altering progenitor specification as a parallel means of controlling IPC production. Shh signaling partly mediates this effect via Gli3R activity, which when absent or reduced, can significantly impair specification of neocortical progenitors into distinct excitatory neuron subtypes (Hasenpusch-Theil et al., 2018; Wang et al., 2011; Yabut et al., 2015). Supporting these findings, we found upregulated expression of known Gli3R gene targets in the E12.5 Sufu-cKO neocortex, such as FGF15, in the progenitor-rich VZ/SVZ where Gli3 is typically highly expressed particularly in RGCs (Hasenpusch-Theil et al., 2015; Kim et al., 2018; Pollen et al., 2014; Rash and Grove, 2007). Specifically, we found an increase in SHH signaling activation (as detected through an increase in Ptch1 expression) correlating with an increase in FGF15 expression in RGCs lining the VZ. However, despite a visibly uniform increase in ectopic FGF15 expression in the E12.5 Sufu-cKO neocortex, we previously observed that RGCs lining the VZ exhibited variable proliferation rates (Yabut et al., 2015). Rather, we consistently found that the increase in FGF15 levels, in the Sufu-cKO embryonic neocortex or when added in cultured forebrain slices, inhibited the specification of RGCs into bona fide pallial IPCs since they begin to ectopically express subpallial-specific markers. Altogether, our findings expand evidence of the importance of modulating Shh signaling in RGCs, particularly via Gli3 activity, in ensuring lineage progression towards the specification of IPCs into distinct excitatory neuron subtypes.

### SHH signaling requires FGF signaling to alter neuronal lineage progression of neocortical progenitors

Our findings revealed that SHH signaling activated FGF signaling leading to disruption of neocortical progenitor specification. Particularly, an increase in FGF15 expression was sufficient to drive these defects. Previous studies showed that FGF15 functions to control progenitor proliferation and differentiation (Borello et al., 2008; Wilson et al., 2012). Supporting a role for FGF15 in proliferation, we found that increased FGF15 expression correlated with the elongation of the E12.5 Sufu-cKO neocortex. However, as mentioned, FGF15 upregulation did not uniformly correlate with proliferation defects. Supporting this, we found variable levels of activation in MAPK signaling, the downstream intracellular FGF signaling effector. MAPK signaling activity was distinctly higher in dorsal and dorsolateral regions of the E12.5 Sufu-cKO neocortex, but not in dorsomedial regions where FGF15 was also ectopically expressed. Additionally, clonally related progenitors in the rostral neocortical regions of the E12.5 Sufu-cKO neocortex appeared to be uniquely affected. Columnar patterns in the expression levels of Pax6, Olig2, and MAPK activity levels are apparent. These observations suggest that FGF15 differentially affects molecularly distinct neocortical progenitors, which may be predicted by their spatial and temporal localization in the embryonic neocortex. These varying effects may lead to changes in cell cycle length, neuronal differentiation, or their specification programs.

While it may be logical to assume that FGF15 will predictably bind to progenitors expressing its cognate receptor, FGF receptor 4 (FGFR4), FGFR4 has not been detected in the E12.5 neocortex (Borello et al., 2008; Harmer et al., 2004; Tole et al., 2006; Zhang et al., 2006). On the other hand, FGFR1-3 are expressed in varying levels and spatial domains across the E12.5 neocortical wall and may be sensitive to high levels of FGF15 (Borello et al., 2008; Tole et al., 2006). Alternatively, the global changes in the molecular make-up of the E12.5 Sufu-cKO neurogenic niche may have facilitated unique FGF15 interactions. This may explain why Sufu-cKO mice do not completely phenocopy mouse mutants carrying conditional Gli3R mutations (Hasenpusch-Theil et al., 2018; Wang et al., 2011). Altogether, these findings underscore the complexity of FGF15 function in the developing neocortex. Future studies require further elucidation of the molecular properties of FGF15-responsive neocortical progenitors and the signaling pathways transduced to elicit changes in progenitor behavior.

## Conclusions

Along with the expansion and diversification of neocortical neuron subtypes in the developing brain, anomalies in the production of diverse excitatory neurons underlie a number of neuropsychiatric and neurodevelopmental disorders. The activation of FGF and MAPK signaling cascades, in response to SHH signaling activation, indicate important potential implications of uncontrolled SHH signaling in these disorders. For instance, it is evident that abnormal numbers and specification of neuronal subtypes lead to aberrant circuits in autism spectrum disorders (ASD) (Kaushik and Zarbalis, 2016). High serum levels of SHH and deregulated FGF signaling activity at developmental stages has been implicated in these defects (Al-Ayadhi, 2012; Halepoto et al., 2015; Rubenstein, 2011; Vaccarino et al., 2009). Activation of SHH signaling at early stages of corticogenesis, consequently driving FGF signaling, may profoundly alter the molecular landscape of neocortical progenitors and their progenies. Thus, further investigation of how pathogenic SHH and FGF signaling converge to produce abnormal neuronal subtypes, and drive abnormal neocortical circuitry, could lay the foundation towards detecting, treating, or even reversing the neocortical abnormalities present in neurodevelopmental disorders.

## EXPERIMENTAL PROCEDURES

### Animals

Mice carrying the floxed Sufu allele (Sufu^fl^) were kindly provided by Dr. Chi-Chung Hui (University of Toronto) and were genotyped as described elsewhere (Pospisilik et al., 2010). *Emx1-cre* (Stock #05628), *Rosa-AI14* (Stock #007908), *SmoM2* (Stock #005130) mice were obtained from Jackson Laboratories (Bar Harbor, ME, USA). Mice designated as controls did not carry the *Cre* transgene and may have either one of the following genotypes: *Sufu^fl/+^* or *Sufu^fl/fl^*. All mouse lines were maintained in mixed strains, and analysis included male and female pups from each age group, although sex differences were not included in data reporting. All animal protocols were in accordance to the National Institute of Health regulations and approved by the UCSF Institutional Animal Care and Use Committee (IACUC).

### RNA-Seq and Analysis

The dorsal forebrain was dissected from E12.5 control and Sufu-cKO littermates (n=4 per group). Total RNA was extracted using Qiagen RNEasy Mini Kit (Qiagen) and prepared for RNAseq. RNASeq was conducted by the UCSF Functional Genomics Core. Barcoded sequencing libraries were generated using the Truseq Stranded mRNA Library Prep Kit (Illumina). Single-end 50-bp reads were sequenced on the HiSeq4000 (Illumina). Sequencing yielded ∼343 million read with an average read depth of 42.9 million reads/sample. Reads were then aligned using STAR_2.4.2a to the mouse genome (Ensembl Mouse GRCm38.78) and those that mapped uniquely to known mRNAs were used to assess differential expression (DE). Final quantification and statistical testing of differentially expressed (adjusted *P* < 0.05) genes were performed using DESeq2. Gene set enrichment and pathway analysis was conducted using The DAVID Gene Functional Classification Tool http://david.abcc.ncifcrf.gov (Huang et al., 2007). Heatmaps represent transformed FPKM values (Transform 1+ Log2(Y)) and plotted using Prism 8.1 (GraphPad). Filtering was applied for GO enrichment analysis by excluding DE genes with very low normalized read counts (FPKM <100) in both control and mutant samples.

### Immunohistochemistry

Perfusion, dissection, and immunofluorescence staining were conducted according to standard protocols as previously described (Siegenthaler et al., 2009). Briefly, embryonic brain tissues were fixed by direct immersion in 4% paraformaldehyde (PFA) and postnatal brains fixed by intracardial perfusion followed by 2 h post-fixation. Cryostat sections were air dried and rinsed 3× in PBS plus 0.2%Triton before blocking for 1 h in 10% normal lamb serum diluted in PBS with 0.2% Triton to prevent nonspecific binding. A heat-induced antigen retrieval protocol was performed on selective immunohistochemistry using 10 μM Citric Acid at pH 6.0. Primary antibodies were diluted in 10% serum diluted in PBS with 0.2% Triton containing 40,6-diamidino-2-phenylindole (DAPI); sections were incubated in primary antibody overnight at room temperature. The following antibodies were used: rabbit anti-Tbr2 (1:500 dilution; Abcam (#ab23345), Cambridge, UK), rabbit anti-GSX2 (1:250 dilution; gift from Kenneth Campbell (Toresson et al., 2000)) mouse anti-Olig2 (1:250 dilution; Millipore (#MABN50), Billerica, MA, USA), phosphor-Erk1/2 (1:100 dilution, Cell Signaling (#4370)). To detect primary antibodies, we used species-specific Alexa Fluor-conjugated secondary antibodies (1:500; Invitrogen) in 1X PBS-T for 1 h at room temperature, washed with 1X PBS, and coverslipped with Fluoromount-G (SouthernBiotech).

### *In Situ* Hybridization

*Lhx2, CoupTF2 in situ* hybridization (ISH) was conducted using RNA probes kindly provided by Professor John Rubenstein (University of California San Francisco). Dlx1 and Dbx1 riboprobles were generated using primer sequences published by the Allen Brain Atlas *ISH* Database (http://developingmouse.brain-map.org/) with SP6 and T7 promoter binding sequences included in 5’ ends. Target gene cDNA was amplified from pooled cDNA reactions made from mouse brain total RNA were used as a template source. DIG-labeled RNA probes were generated using the DIG RNA Labeling Kit SP6/T7 (Sigma-Aldrich cat. # 11175025910) according to manufacturer’s protocols. DIG-labeled RNA probes were diluted in hybridization buffer (50% formamide, 5 x SSC, 0.3 mg/ml tRNA, 100 µl/ml Heparin, 1x Denhardt’s solution, 0.1% Tween 20, 0.1% CHAPS, 5 mM EDTA) and added to RNase-free cryosections for incubation in a humidified chamber at 65°C for 16-20h. Sections were washed in 0.2 x SSC (Ambion AM9770) at 65°C followed by PBST at room temperature. Tissue sections were incubated in alkaline phosphatase-conjugated anti-DIG antibody (1:1500, Roche Applied Sciences 11093274910) for 16-20h incubation at room temperature and colorimetric signals were detected using Nitro blue tetrazolium chloride and 5-bromo-4-chloro-3-indolyl phosphase (NBT-BCIP; Roche Applied Sciences 11383221001). RNAScope ISH was conducted for FGF15 and Ptch1. RNAscope probes Mm-Ptch1 (Cat No. 402811) and Mm-FGF15 (Cat No. 412811) were designed commercially by the manufacturer (Advanced Cell Diagnostics, Inc.). RNAScope Assay was performed using the RNAscope Multiplex Fluorescent Reagent Kit V2 according to manufacturer’s instructions. Detection of the probe was done with Opal 570 or Opal 520 reagent (Perkin Elmer).

### Forebrain Organotypic Slice Culture

Whole brains from E12.5 wildtype CD-1 mice were carefully dissected and placed in ice-cold Hanks Balanced Salt Solution (HBSS; Invitrogen). Brains were embedded in 4% Low Melting Point Agarose (Nueve)/HBSS mix and allowed to solidify on ice. Embedded brains were sliced using a VT1000S vibratome (Leica) into 400 μm thick slices and placed in Recovery Media (MEM (Invitrogen) with Glutamax (Invitrogen) and Pennicillin/Streptomycin (Invitrogen)). Slices were transferred into uncoated Millicell-CM membrane inserts (EMD-Millipore) in 6-well plates (BD Biosciences) and cultured in Neurobasal (Invitrogen) supplemented with Glutamax (Invitrogen), Pennicillin/Streptomycin (Invitrogen), B-27 (Invitrogen) and N2 (Invitrogen) at 37°C, 5% CO_2_, and 100% humidity. After 2 days in vitro (DIV), cell culture media were aspirated, and slices were washed in 1X PBS, fixed in cold 4% PFA for 30 minutes, cryoprotected in 30% sucrose, and embedded in OCT. Slices were cryosectioned into 20 μm thick coronal sections and stored at −80°C until used for immunofluorescence analysis as described above. Treatment (as described in text) of organotypic slices were conducted 2-3 hours after initial plating and incubation of slices with the following concentrations: 100 ng/ml recombinant FGF15 (Prospec Bio, #CYT-027), 200 ng/ml recombinant SHH (GenScript, #Z03050-50), and 5 uM cyclopamine (Toronto Research Chemicals, #C988400). Following treatments, slice cultures were incubated for 2 days and processed as described above.

### Image Analysis and Acquisition

Images were acquired using a Nikon E600 microscope equipped with a QCapture Pro camera (QImaging), Zeiss Axioscan Z.1 (Zeiss, Thornwood, NY, USA) using the Zen 2 blue edition software (Zeiss, Thornwood, NY, USA), or the Nikon Ti inverted microscope with CSU-W1 large field of view confocal and Andor Zyla 4.2 sCMOS camera. All images were imported in tiff or jpeg format. Brightness, contrast, and background were adjusted equally for the entire image between controls and mutant using the “Brightness/Contrast” and “Levels” function from “Image/Adjustment” options in Adobe Photoshop or NIH ImageJ without any further modification. NIH Image J was used to threshold background levels between controls and mutant tissues to quantify fluorescence labeling. For Phospho-Erk1/2 quantification, the total area with positive Phospho-Erk1/2 labeling were measured, which began in the pallial-subpallial boundary in the controls and extended dorsally in Sufu-cKO neocortex, for each hemisphere across the anterior to posterior axis. One forebrain section in each representative anterior to posterior region were measured (**Figure 5C**) from both hemispheres were averaged. All analyses were conducted in at least 2-3 20 μm thick sections that were histologically matched at the rostral-caudal level between genotypes.

### Statistics

Prism 8.1 (GraphPad) was used for statistical analysis. Two sample experiments were analyzed by Student’s *t* test and experiments with more than two parameters were analyzed by ANOVA. In two- or three-way ANOVA, when interactions were found, follow up analysis were conducted for the relevant variables using Holm-Sidak’s multiple comparisons test. All experiments were conducted in triplicate with a sample size of n = 3−6 embryos/animals/slices per genotype. *P* value ≤0.05 were considered statistically significant. Graphs display the mean ± standard error of the mean (SEM). Statistical values are shown in Supplemental Table 4.

## Acknowledgements

We thank members of the Pleasure Lab for helpful discussions, Dr. Kenneth Campbell for the Gsx2 antibody, and DeLaine Larsen and Kari Harrington at the University of California San Francisco Nikon Imaging Center for assistance with imaging. This work was supported by the NIH R01s MH077694 and NS118995 (S.J.P.), NIH/NCI K01CA201068 (O.R.Y.), and KNRF 2019M3A9H1103702 (K.Y).

## Extended Data Figures

**Extended Data Figure 1-1:**
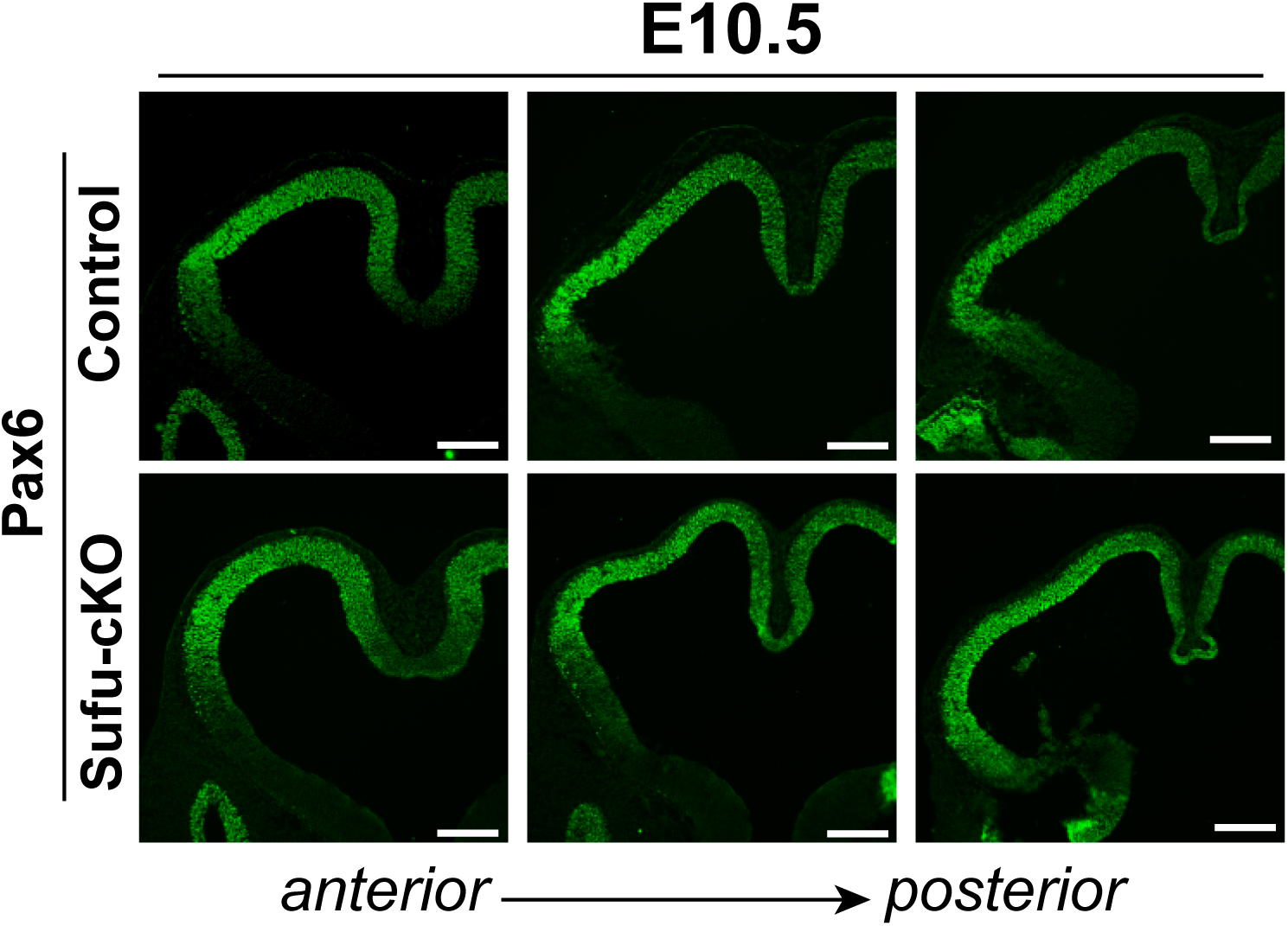
Immunofluorescence staining of the E10.5 forebrain for the pallial marker, Pax6, showed exclusive expression of Pax6 in dorsal forebrain regions in both control and Sufu-cKO mice. Scale bar = 200 μm.

**Extended Data Figure 3-1:**
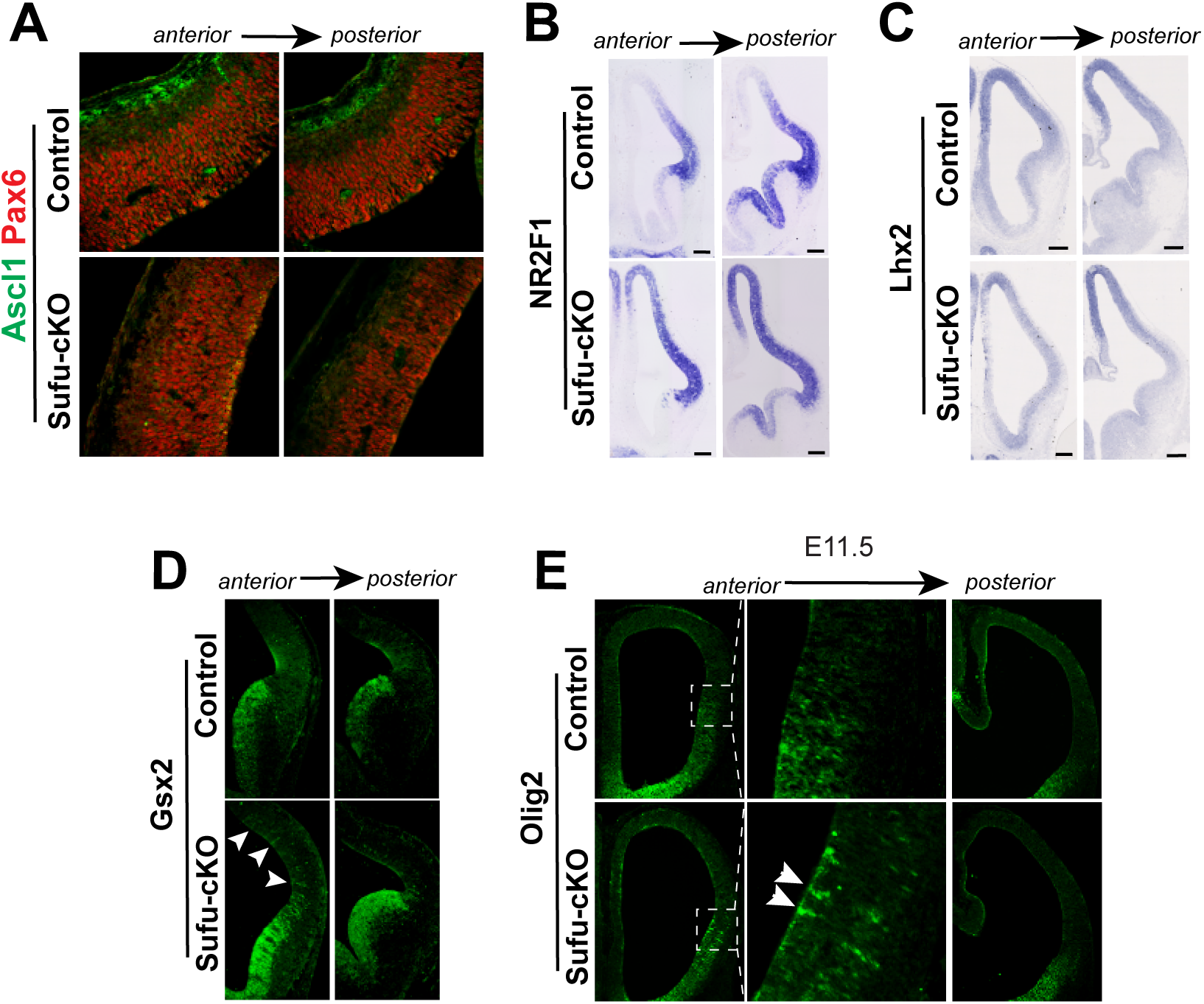
Increased expression of ventral progenitor markers in neocortical progenitors of E12.5 Sufu-cKO embryos. (A) Immunofluorescence staining for Ascl1 and Pax6 on representative coronal sections of E12.5 control and Sufu-cKO neocortex showed very low levels of Ascl1 protein expression in the E12.5 neocortex with either genotypes. (B, C) In situ hybridization using riboprobes for NRF2F1 (B) and Lhx2 (C) were conducted on representative coronal sections of E12.5 control and Sufu-cKO neocortex showed that these pallial-specific markers were comparably expressed in both control and mutant mice. Scale bar = 200 μm. (D) Immunofluorescence staining for the subpallial-specific marker, Gsx2, on representative coronal sections of E12.5 control and Sufu-cKO neocortex. Gsx2 was detected in the anterior regions of the E12.5 Sufu-cKO neocortex and showed a columnar pattern of expression. (E) Immunofluorescence staining for the subpallial-specific marker, Olig2, on representative coronal sections of E11.5 control and Sufu-cKO neocortex showed irregularities in Olig2 expression in the Sufu-cKO neocortex, particularly near the pallial-subpallial boundary.

**Extended Data Figure 3-2:**
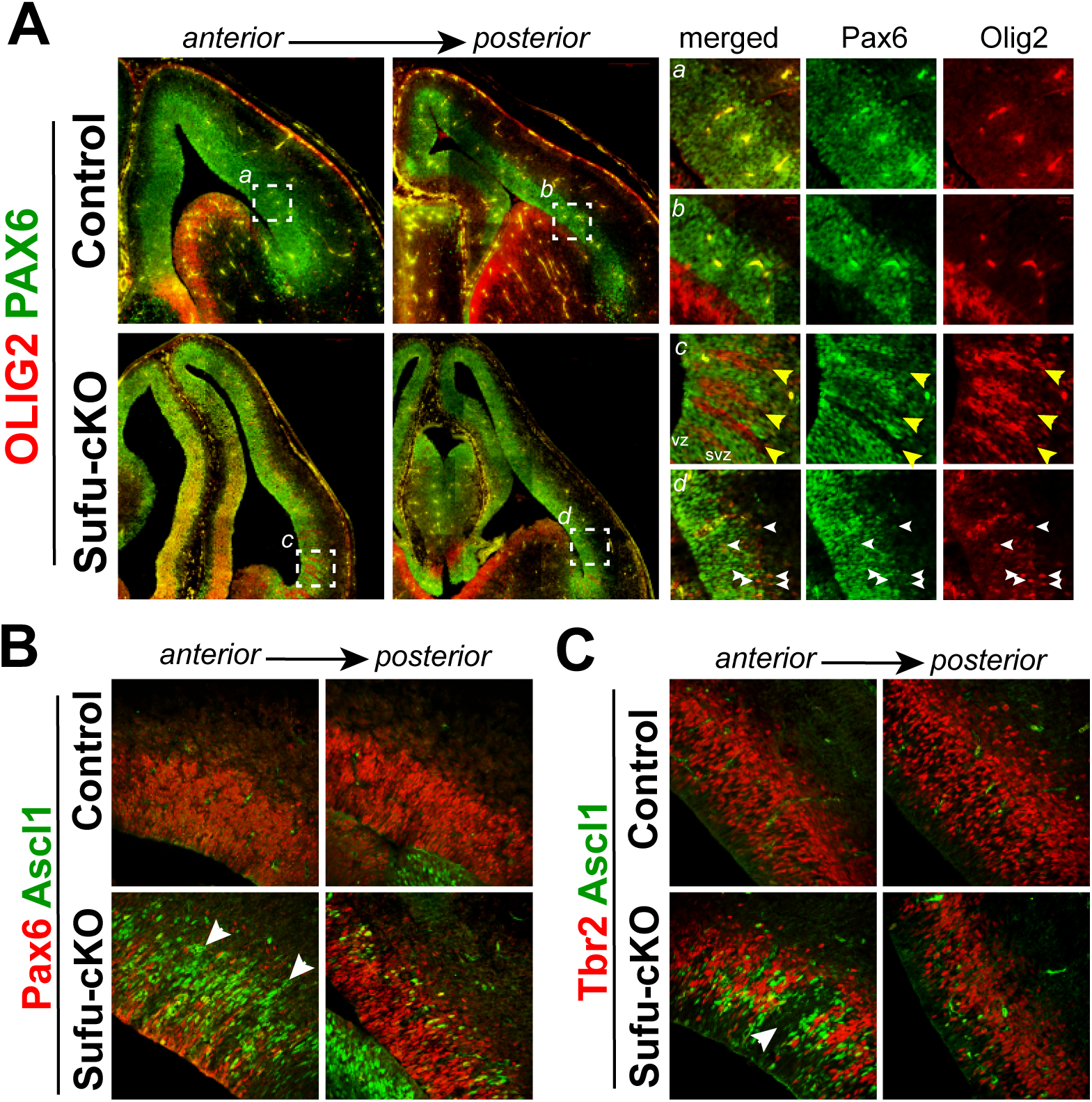
Ectopic expression of Olig2 precedes Ascl1 expression. (A) Immunofluorescence staining for Olig2 and Pax6 on representative coronal sections of E14.5 control and Sufu-cKO neocortex. Ectopic expression of Olig2 was detected across the A-P axis, with a columnar pattern of expression prevalent in anterior regions of the E14.5 Sufu-cKO neocortex. (B, C) Immunofluorescence staining for Ascl1 and Pax6 (B) or Tbr2 (C) on representative coronal sections of E14.5 control and Sufu-cKO neocortex.Ectopic expression of Ascl1 was detected across the A-P axis of the E14.5 Sufu-cKO neocortex. However, more Ascl1+ cells were detected in anterior regions.

**Extended Data Figure 4-1:**
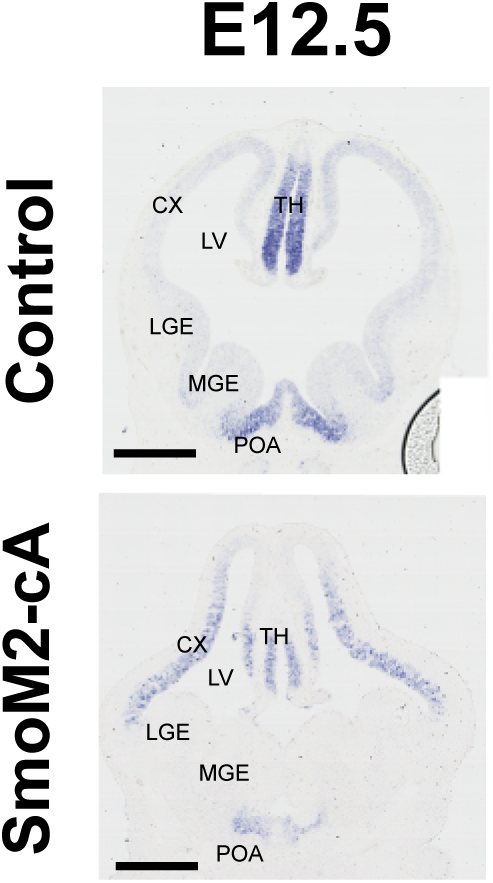
Constitutive activation of Shh signaling drives ectopic Fgf15 expression in the embryonic neocortex. *In situ* hybridization using Fgf15-specific riboprobes on E12.5 embryos carrying the constitutively-active Smoothened allele in neocortical progenitors (Emx1-Cre;SmoM2 or SmoM2-cA), showed upregulation of Fgf15 expression.

## BIBLIOGRAPHY

Al-Ayadhi LY. 2012. Relationship Between Sonic Hedgehog Protein, Brain-Derived Neurotrophic Factor and Oxidative Stress in Autism Spectrum Disorders. Neurochem Res 37:394–400. doi:10.1007/s11064-011-0624-x

Beattie R, Hippenmeyer S. 2017. Mechanisms of radial glia progenitor cell lineage progression. FEBS Lett 591:3993–4008. doi:10.1002/1873-3468.12906

Borello U, Cobos I, Long JE, Murre C, Rubenstein JL. 2008. FGF15 promotes neurogenesis and opposes FGF8 function during neocortical development. Neural Dev 3:17. doi:10.1186/1749-8104-3-17

Dave RK, Ellis T, Toumpas MC, Robson JP, Julian E, Adolphe C, Bartlett PF, Cooper HM, Reynolds BA, Wainwright BJ. 2011. Sonic hedgehog and notch signaling can cooperate to regulate neurogenic divisions of neocortical progenitors. PLoS One 6:e14680. doi:10.1371/journal.pone.0014680

Fotaki V, Yu T, Zaki PA, Mason JO, Price DJ. 2006. Abnormal positioning of diencephalic cell types in neocortical tissue in the dorsal telencephalon of mice lacking functional Gli3. J Neurosci 26:9282–92. doi:10.1523/JNEUROSCI.2673-06.2006

Gimeno L, Martinez S. 2007. Expression of chickFgf19 and mouseFgf15 orthologs is regulated in the developing brain byFgf8 andShh. Dev Dyn 236:2285–2297. doi:10.1002/dvdy.21237

Guillemot F, Zimmer C. 2011. From Cradle to Grave: The Multiple Roles of Fibroblast Growth Factors in Neural Development. Neuron 71:574–588. doi:10.1016/J.NEURON.2011.08.002

Halepoto DM, Bashir S, Zeina R, Al-Ayadhi LY. 2015. Correlation Between Hedgehog (Hh) Protein Family and Brain-Derived Neurotrophic Factor (BDNF) in Autism Spectrum Disorder (ASD). J Coll Physicians Surg Pak 25:882–5. doi:12.2015/JCPSP.882885

Harmer NJ, Pellegrini L, Chirgadze D, Fernandez-Recio J, Blundell TL. 2004. The crystal structure of fibroblast growth factor (FGF) 19 reveals novel features of the FGF family and offers a structural basis for its unusual receptor affinity. Biochemistry 43:629–40. doi:10.1021/bi035320k

Hasenpusch-Theil K, Watson JA, Theil T. 2015. Direct Interactions Between Gli3, Wnt8b, and Fgfs Underlie Patterning of the Dorsal Telencephalon. Cereb Cortex 27:bhv291. doi:10.1093/cercor/bhv291

Hasenpusch-Theil K, West S, Kelman A, Kozic Z, Horrocks S, McMahon AP, Price DJ, Mason JO, Theil T. 2018. Gli3 controls the onset of cortical neurogenesis by regulating the radial glial cell cycle through Cdk6 expression. Development 145. doi:10.1242/dev.163147

Hevner RF. 2019. Intermediate progenitors and Tbr2 in cortical development. J Anat 235:616–625. doi:10.1111/joa.12939

Huang DW, Sherman BT, Tan Q, Collins JR, Alvord WG, Roayaei J, Stephens R, Baseler MW, Lane HC, Lempicki RA. 2007. The DAVID Gene Functional Classification Tool: a novel biological module-centric algorithm to functionally analyze large gene lists. Genome Biol 8:R183. doi:10.1186/gb-2007-8-9-r183

Kaushik G, Zarbalis KS. 2016. Prenatal Neurogenesis in Autism Spectrum Disorders. Front Chem 4:12. doi:10.3389/fchem.2016.00012

Kim JJ, Jiwani T, Erwood S, Loree J, Rosenblum ND. 2018. Suppressor of fused controls cerebellar neuronal differentiation in a manner modulated by GLI3 repressor and Fgf15. Dev Dyn 247:156–169. doi:10.1002/dvdy.24526

Komada M, Saitsu H, Kinboshi M, Miura T, Shiota K, Ishibashi M. 2008a. Hedgehog signaling is involved in development of the neocortex. Development 135:2717–27. doi:10.1242/dev.015891

Komada M, Saitsu H, Shiota K, Ishibashi M. 2008b. Expression of Fgf15 is regulated by both activator and repressor forms of Gli2 in vitro. Biochem Biophys Res Commun 369:350–356. doi:10.1016/J.BBRC.2008.02.015

Long F, Zhang XM, Karp S, Yang Y, McMahon AP. 2001. Genetic manipulation of hedgehog signaling in the endochondral skeleton reveals a direct role in the regulation of chondrocyte proliferation. Development 128:5099–5108.

Miyoshi G, Butt SJB, Takebayashi H, Fishell G. 2007. Physiologically distinct temporal cohorts of cortical interneurons arise from telencephalic Olig2-expressing precursors. J Neurosci 27:7786–98. doi:10.1523/JNEUROSCI.1807-07.2007

Palma V, Ruiz i Altaba A. 2004. Hedgehog-GLI signaling regulates the behavior of cells with stem cell properties in the developing neocortex. Development 131:337–45. doi:10.1242/dev.00930

Petryniak MA, Potter GB, Rowitch DH, Rubenstein JLR. 2007. Dlx1 and Dlx2 control neuronal versus oligodendroglial cell fate acquisition in the developing forebrain. Neuron 55:417–33. doi:10.1016/j.neuron.2007.06.036

Pollen AA, Nowakowski TJ, Shuga J, Wang X, Leyrat AA, Lui JH, Li N, Szpankowski L, Fowler B, Chen P, Ramalingam N, Sun G, Thu M, Norris M, Lebofsky R, Toppani D, Kemp DW, Wong M, Clerkson B, Jones BN, Wu S, Knutsson L, Alvarado B, Wang J, Weaver LS, May AP, Jones RC, Unger MA, Kriegstein AR, West JAA. 2014. Low-coverage single-cell mRNA sequencing reveals cellular heterogeneity and activated signaling pathways in developing cerebral cortex. Nat Biotechnol 32:1053–1058. doi:10.1038/nbt.2967

Rash BG, Grove EA. 2007. Patterning the dorsal telencephalon: a role for sonic hedgehog? J Neurosci 27:11595–603. doi:10.1523/JNEUROSCI.3204-07.2007

Rubenstein JLR. 2011. Annual Research Review: Development of the cerebral cortex: implications for neurodevelopmental disorders. J Child Psychol Psychiatry 52:339–355. doi:10.1111/j.1469-7610.2010.02307.x

Shikata Y, Okada T, Hashimoto M, Ellis T, Matsumaru D, Shiroishi T, Ogawa M, Wainwright B, Motoyama J. 2011. Ptch1-mediated dosage-dependent action of Shh signaling regulates neural progenitor development at late gestational stages. Dev Biol 349:147–59. doi:10.1016/j.ydbio.2010.10.014

Siegenthaler JA, Ashique AM, Zarbalis K, Patterson KP, Hecht JH, Kane MA, Folias AE, Choe Y, May SR, Kume T, Napoli JL, Peterson AS, Pleasure SJ. 2009. Retinoic acid from the meninges regulates cortical neuron generation. Cell 139:597–609. doi:10.1016/j.cell.2009.10.004

Sohal VS, Rubenstein JLR. 2019. Excitation-inhibition balance as a framework for investigating mechanisms in neuropsychiatric disorders. Mol Psychiatry 24:1248–1257. doi:10.1038/s41380-019-0426-0

Tole S, Gutin G, Bhatnagar L, Remedios R, Hébert JM. 2006. Development of midline cell types and commissural axon tracts requires Fgfr1 in the cerebrum. Dev Biol 289:141–51. doi:10.1016/j.ydbio.2005.10.020

Toresson H, Potter SS, Campbell K. 2000. Genetic control of dorsal-ventral identity in the telencephalon: opposing roles for Pax6 and Gsh2. Development 127:4361–71.

Vaccarino FM, Grigorenko EL, Smith KM, Stevens HE. 2009. Regulation of Cerebral Cortical Size and Neuron Number by Fibroblast Growth Factors: Implications for Autism. J Autism Dev Disord 39:511–520. doi:10.1007/s10803-008-0653-8

Wang H, Ge G, Uchida Y, Luu B, Ahn S. 2011. Gli3 is required for maintenance and fate specification of cortical progenitors. J Neurosci 31:6440–8. doi:10.1523/JNEUROSCI.4892-10.2011

Wang L, Hou S, Han Y-G. 2016. Hedgehog signaling promotes basal progenitor expansion and the growth and folding of the neocortex. Nat Neurosci 19:888–896. doi:10.1038/nn.4307

Wang Y, Kim E, Wang X, Novitch BG, Yoshikawa K, Chang L-S, Zhu Y. 2012. ERK Inhibition Rescues Defects in Fate Specification of Nf1-Deficient Neural Progenitors and Brain Abnormalities. Cell 150:816–830. doi:10.1016/j.cell.2012.06.034

Wilson SL, Wilson JP, Wang C, Wang B, McConnell SK. 2012. Primary cilia and Gli3 activity regulate cerebral cortical size. Dev Neurobiol 72:1196–212. doi:10.1002/dneu.20985

Yabut O, Ng H, Fernandez G, Yoon K, Kuhn J, Pleasure S. 2016. Loss of Suppressor of Fused in Mid-Corticogenesis Leads to the Expansion of Intermediate Progenitors. J Dev Biol 4:29. doi:10.3390/jdb4040029

Yabut OROR, Fernandez G, Huynh T, Yoon K, Pleasure SJSJ. 2015. Suppressor of Fused Is Critical for Maintenance of Neuronal Progenitor Identity during Corticogenesis. Cell Rep 12:2021–34. doi:10.1016/j.celrep.2015.08.031

Ypsilanti AR, Rubenstein JLR. 2016. Transcriptional and epigenetic mechanisms of early cortical development: An examination of how Pax6 coordinates cortical development. J Comp Neurol 524:609–629. doi:10.1002/cne.23866

Zhang X, Ibrahimi OA, Olsen SK, Umemori H, Mohammadi M, Ornitz DM. 2006. Receptor specificity of the fibroblast growth factor family. The complete mammalian FGF family. J Biol Chem 281:15694–700. doi:10.1074/jbc.M601252200

